# Potential short-term earthquake forecasting by farm-animal monitoring

**DOI:** 10.1101/2020.01.19.911313

**Authors:** Martin Wikelski, Uschi Mueller, Paola Scocco, Andrea Catorci, Lev Desinov, Mikhail Belyaev, Daniel Keim, Winfried Pohlmeier, Gerhard Fechteler, P. Martin Mai

## Abstract

Whether changes in animal behavior allow for short-term earthquake predictions has been debated for a long time. During the 2016/2017 earthquake sequence in Italy, we instrumentally observed the activity of farm animals (cows, dogs, sheep) close to the epicenter of the devastating magnitude M6.6 Norcia earthquake (Oct-Nov 2016) and over a subsequent longer observation period (Jan-Apr 2017). Relating 5304 (in 2016) and 12948 (in 2017) earthquakes with a wide magnitude range (0.4 ≤ M ≤ 6.6) to continuously measured animal activity, we detected how the animals collectively reacted to earthquakes. We also found consistent anticipatory activity prior to earthquakes during times when the animals were in a stable, but not during their time on a pasture. We detect these anticipatory patterns not only in periods with high, but also in periods of low seismic activity. Earthquake anticipation times (1-20hrs) are negatively correlated with the distance between the farm and earthquake hypocenters. Our study suggests that continuous instrumental monitoring of animal collectives has the potential to provide statistically reliable patterns of pre-seismic activity that could allow for short-term earthquake forecasting.

**One Sentence Summary:** A collective of domestic animals repeatedly showed unusually high activity levels before earthquakes, with anticipation times (1-20h) negatively related to distance from epicenters (5-28km).

## Introduction

Earthquakes are a major threat as they strike unexpectedly in exact space and time, causing large economic and societal losses [1–5]. While the general location, time period, and expected magnitude range can be statistically forecasted for earthquakes in well-instrumented regions, accurate short-term predictions are considered impossible [6–8]. To prepare for earthquakes and their consequences, probabilistic seismic hazard assessment is used to estimate possible shaking levels for future earthquakes. Earthquake early warning systems provide automated short-notice local warnings to agencies and infrastructural systems about the imminent danger of strong shaking [1, 9]. However, reliable technical warning systems that anticipate the location, magnitude and timing of an earthquake within minutes to hours do not exist [8, 10].

Since ancient times abnormal animal behavior prior to earthquakes or volcanic eruptions has been described, i.e., some animals show “aberrant” or “strange” behavior in anticipation of natural disasters [2, 11, 12]. Most famously, the 1975 Haicheng earthquake (magnitude M 7.2) in China, was anticipated based on human observations of animal behavior, such as snakes or rats leaving their burrows in winter [13]. Similar observations are rare [14], but recently evidence accumulated that animals in earthquake areas may show aberrant behavior [15–22]. Nevertheless, a recent review [23] points out the sparsity of data and need for testable quantitative measures on animal-anticipated earthquake occurrence.

Assuming that measurable physical precursors for earthquakes exist, three conditions must be met for animal behavior to be possibly useful for short-term earthquake forecasting [24]: i) The precursors must be perceived by animals; ii) Animals must respond to precursors by showing measurable, quantifiable, and testable behavioral patterns; iii) These behavioral patterns must be detected and clearly distinguished against the background of regular behavior. In many reports on anticipatory animal behavior, these three conditions have been met only partially [25–29].

More recently, several approaches proposed to quantify animal behavior [11, 24–27, 30–35] in accordance with the abovementioned conditions. Amongst others, the use of camera traps for birds and mammals [11, 24–27, 30–35] and the use of locomotor sensors for mice [11, 24–27, 30–35] have shown potential to be useful to detect behavioral changes in animal behavior prior to earthquakes.

In our study we used bio-logging techniques [36], enabling remote, continuous observation of animals in unprecedented detail, particularly through continuous 3D-accelerometer data [37] [38]. Moreover, recent advances in understanding animal behavior show that collectives of animals can have sensing abilities that outperform individuals [39]. Collectives are defined here as interacting groups of animals, either within or between species. Thus, one may speculate that some animal collectives are able to detect and process physical signals [40] for which currently no engineered recording devices exist. Correspondingly, our study is not only aimed at providing evidence for unusual animal behavior prior to earthquakes (as proposed in [41]), but also at verifying that animals continuously respond to changes in potential precursors of earthquakes. For this purpose, we measured the activity (overall dynamic body acceleration, ODBA) of multiple cows, dogs and sheep at a farm nearby the hypocenter of the M6.6 Norcia (Italy) earthquake and analyzed them in the context of the ongoing seismicity. We distinguish three time periods: (a) the Oct-Nov 2016 period shortly before and after the M6.6 Norcia shock where the animals were in the stable; (b) the Jan-Mar 2017 period of lower earthquake activity, where the animals were also in the stable; (c) the Mar-Apr 2017 period, where the animals were on a pasture.

A number of possible precursory processes and associated physical signal have been suggested in the literature [11, 24–27, 30–35], but there is no consensus on which of them may explain changes in animal behavior. Our considerations are based on the assumption that a diffusive process, possibly related to slow deformation processes in the rock volume near the hypocentral region of the ensuing earthquake, generates and emanates a physical measurable precursory signal. However, we refrain from speculating about the details of the potential mechanisms of this diffusive process. The goal of this study is to measure and analyze the anticipatory patterns without relying on assumptions about a potential mechanism.

## Results

### Daily patterns in animal behavior

To identify unusual animal behavior one first has to identify and quantify the (daily) normal activity patterns. Therefore, we examined statistically robust daily activity patterns for three animal species (cows, dogs and sheep); these are then considered in our analysis. For the time periods (a) and (b) when animals were in the stable (Oct-Nov 2016 and Jan-Mar 2017), we find that all three species show high activity during the morning and afternoon, but lower activity during noon and at night. During period (c) on the pasture (Mar-Apr 2017), only cows showed a significantly reduced activity during noon.

### Mutual influence of the animal species

All tagged animals were held on the same farm. To further understand the normal activity patterns of the animals, we studied the mutual influence of the three species on each other. During all three periods, cows and dogs significantly reacted upon each other; only in the period on the pasture (Mar-Apr 2017) we find a strong reaction of sheep on dog activity, because the dogs guarded the sheep on the pasture. The mutual reaction patterns are presented in Figures S7 and S8 in the supplementary materials

### Reactive animal behavior after earthquakes

In addition, we considered the reaction of the three animal species on earthquakes. We detected that the species differ in their sensibility towards earthquakes. Dogs were most sensible, followed by cows, while the sheep’ activity hardly changed. Moreover, also the reactive patterns differ between the species. While dogs became hyperactive as response to earthquakes, cows initially became untypically calm, but then increased their activity in response to the dogs’ activeness. Notably, these reaction patterns were only found during the periods when the animals were in the stable (Oct-Nov 2016 and Jan-Mar 2017), but not during the time on the pasture (Mar-Apr 2017).

### Anticipatory animal behavior prior to earthquakes

Finally, knowing the normal activity patterns of the animal species, we analyzed the potential anticipatory behavior of the animals prior to earthquakes. Based on a threshold approach (see the Methods section), we identified the anticipation times (time difference between unusually high animal activity and the subsequent earthquake) of the animal species. For a slow diffusive process that generates the precursory signal that the animals react to, we expect the anticipation (or warning) times to depend inversely on hypocentral distance of the respective earthquake to the farm. The further away the earthquake, the shorter the animal warning time. We detect this relationship for both periods in which the animals were in the stable, for the Oct-Nov 2016 period with strong earthquake activity (Fig. 1A), as well as for the Jan-Apr 2017 period with lower earthquake activity (Fig. 1B). For the period when the animals freely roamed on the pasture (Mar-Apr 2017), no significant relationship between anticipation times and hypocentral distances could be found (Fig. 1C). Interestingly, aggregating the information from the three animal collectives helps to identify and establish the statistical significance of this inverse relationship (reflected by the negative slope in Fig. 1A). The pattern was insignificant when considering the information on the individual species (Fig. S9). This indicates that the aggregation is likely to reduce background noise.

**Fig. 1.**
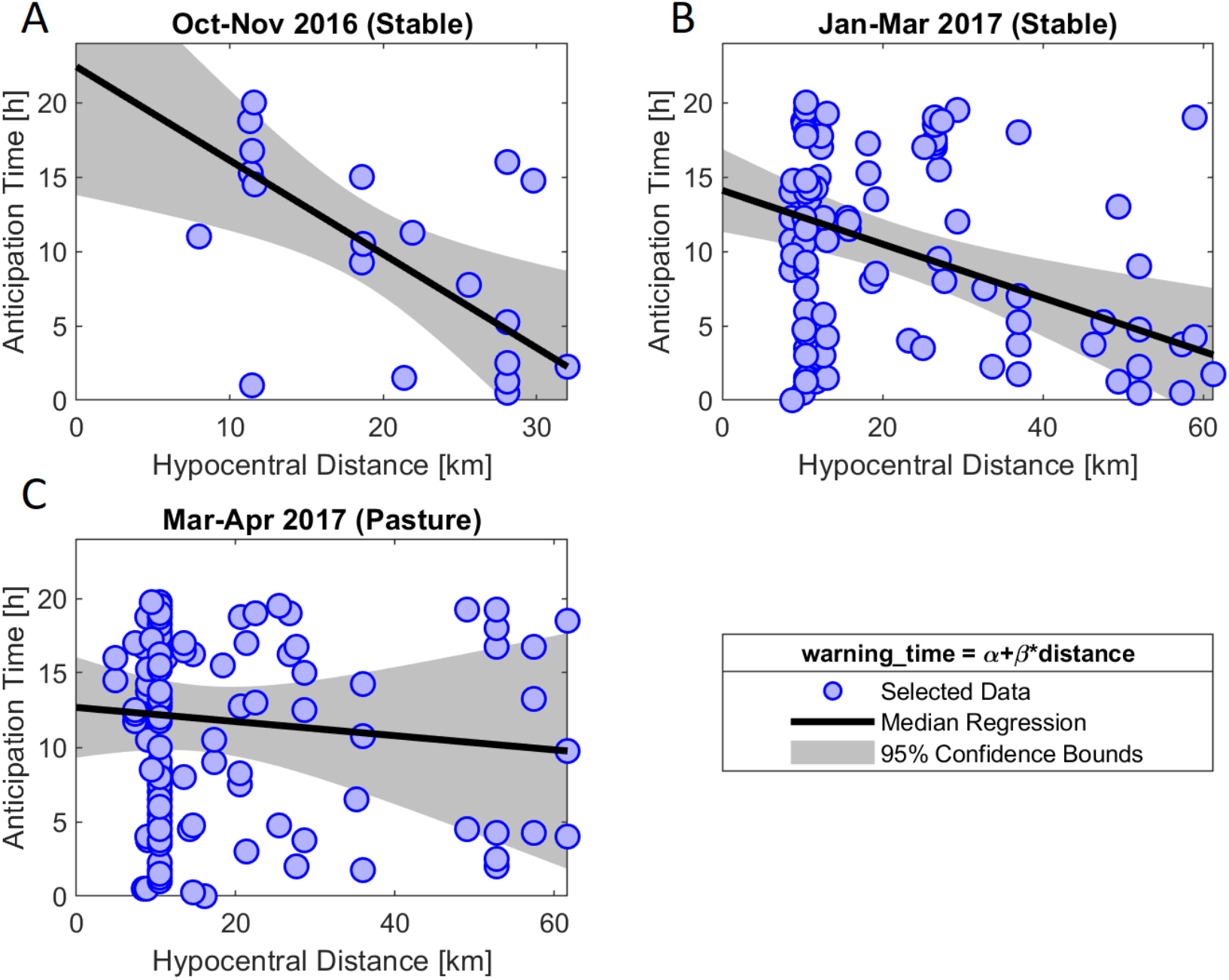
Warning time before an earthquake event (time difference between earthquake event and increased animal activity) against the distance between the respective hypocenter and the farm. Assuming that physical precursors of earthquakes diffuse slowly from the respective hypocenter, we expect a relationship with negative slope when plotting anticipation time against hypocentral distance. A) Oct-Nov 2016 period B) Jan-Mar 2017 period. Same pattern as in A), both periods when the animals were in a stable show a significantly negative relationship. C) Mar-Apr 2017 period. For this period, where the animals were on the pasture, we do not find a significant relationship.

## Discussion

Our observational study systematically quantifies the inter-relationships of complete time series of animal activity and earthquake occurrence (including large mainshocks and their aftershocks), as well as during a periods of lower EQ activity [1, 42, 43]. Our animal activity data are based on proven behavioral surveillance methods that quantify activity patterns of a collective of domestic animal species [36–38].

The usual behavior of the animals is subject to a strong daily pattern and mutual interactions between the three observed species. Moreover, reactive patterns to seismic activity are observed during the periods when the animals were in a stable (Oct-Now 2016 and Jan-Mar 2017).

The analysis of the anticipatory animal behavior provides evidence that the animals are steadily influenced by changes in the physical precursor of seismic events. Notably, the detection of the anticipatory patterns does not rely on the occurrence of a few strong and rare earthquakes, but it is also obtained for periods with medium-size earthquakes. This eases the detection of larger earthquakes against the background of noise in the animals’ activity.

Both, the reactive and the anticipatory behavior of the animals was significant for the periods when the animals were in a stable (Oct-Now 2016 and Jan-Mar 2017), but not when they were on a pasture (Mar-Apr 2017). This implies that the animals are more sensitive in closed buildings. However, our conjecture cannot rule out the possibility that there may exist simple seasonal differences in behavior. But as the patterns are equivalent for the Oct-Nov 2016 and the Jan-Mar 2017 period, both with the animals held in a stable, the distinction between periods in a stable and periods on a pasture seems to be the decisive and most reasonable one.

Overall, the continuous monitoring of animal behavior over longer time spans at high temporal sampling (on the order of minutes) in the controlled setting of a stable allow us to identify the anticipatory behavior, irrespective of the occurrence of large earthquakes.

Our finding that the anticipation time depends inversely on hypocentral distance is consistent with a (slow) diffusion-like mechanism, originating in the rock volume around the earthquake nucleation region at depth (about 5-18 km, see Figure S5c). This process seems qualitatively related to the pre-slip model for earthquake nucleation [44]. The results indicate that the anticipation time might span up to 15-25 hours. Given that the maximum anticipation time and slope of the anticipation time - hypocentral distance - plot can be estimated more precisely in further experiments, these parameters might be useful to identify the actual precursors that the animals react to. One mechanism consistent with our observations was proposed by Freund and Stolc [45]. The air ionizations at pressurized rock surfaces could slowly diffuse in the air towards the animals that then react towards this novel sensation.

In the experimental setup discussed above, the animal activity at a single farm cannot identify in advance the time and distance of a future earthquake [11]. Either an earthquake will occur at larger distance from the farm, but soon, or it will happen close-by, but not as soon. To estimate both time and location of a future earthquake, we propose an experimental animal-sensor triangulation system to forecast the most likely epicenter of an earthquake (Fig. 2).

**Fig. 2.**
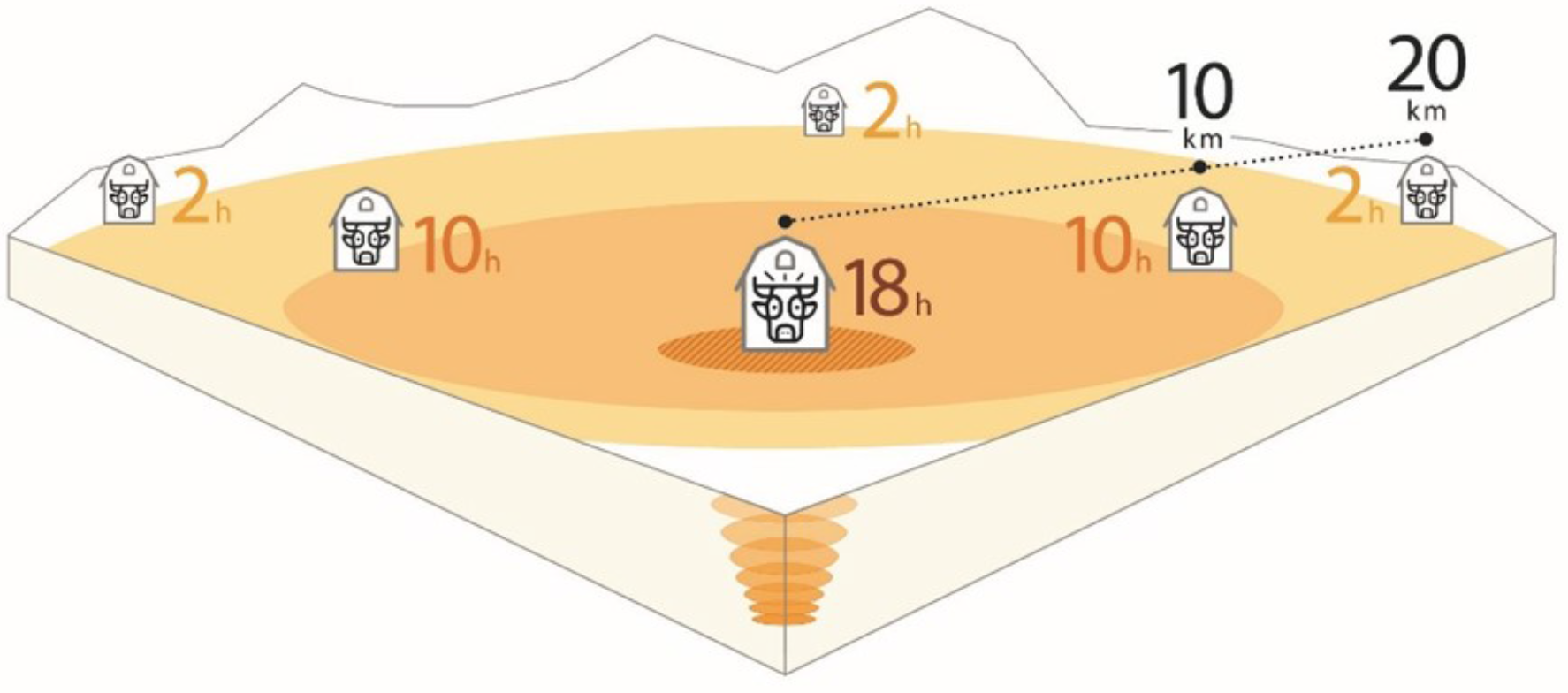
A potential test setup for an earthquake-forecast-by-animals scenario. Based on the information in Fig. 1, one could set up animal activity monitoring sites across a landscape (e.g., the Italian Abruzzo mountains). If there are preparatory processes in the Earth crust indicating an imminent earthquake that could be perceived by animals, the animals on the central farm situated above the hypocenter should show activity ca. 18h ahead of the earthquake, but no other farm animals should show warning signs. Then, 8 hours later, farm animals at 10km distance from the central farm should show warning signs. Another 8 hours later animals at farms 20km away from the central farm should show warning signs. If correct, this would indicate an earthquake is imminent within the next 2 hours. Note that the time constants of 18hrs and 8hrs used in this schematic are indicative examples only. If this hypothesis is tested via experiments and theory involving a physical mechanism, these parameters are expected to change.

All in all, our findings demonstrate that a continuous monitoring of animal collectives at different locations has the potential to detect statistically reliable patterns of pre-seismic activity, that is promising for short-time earthquake forecasting.

## Methods

### Field site selection and tagging of animals

We approached the farm of the Angeli brothers in the village of Capriglia (Fig. S1), and upon consultation with local authorities, received oral permission and the help of the farmers to tag their domestic animals. We chose to tag the animal species and individuals that were selected by the farmers as being potentially sensitive to earthquakes, based on the farmers previous experience. On Oct 28^th^ 2016, we tagged a total of 6 cows, 5 sheep and 2 dogs within 3-30 kilometers of subsequent earthquakes (S1,2), using 54Hz-3D-acceleration loggers to continuously quantify their overall dynamic body acceleration (ODBA), a measure for animal activity [37]. The loggers were synchronized to GPS time immediately before deployment and were set to start recording at 18h UTC on Oct 28^th^, 2016. We left the farm on Oct 28^th^ 2016, at 15h UTC. We then returned to the farm on Nov 18^th^ 2016 to retrieve the tags. Data were downloaded immediately, entered into Movebank [46–48] and visually pre-analyzed.

We returned again to the farm on Jan 3^rd^ 2017 to record additional animal activity data; it turned out that earthquake activity was reduced in that period. We tagged the same individual animals again that were previously tagged, from Jan 17^th^ until Apr 16^th^ 2017. During the winter period (Oct-Mar 11^th^ 2017), the cows were held in a stable, chained to one predefined location as is customary in traditional farms (Fig. S1, S2). The dogs were generally kept inside the house or in the narrow courtyard, from which they could also enter the stables of the cows or sheep. Starting from Mar 11^th^ 2017, the animals were brought to the pastures that surround the farm (Fig. S3) and could roam freely within their large enclosures.

All our experiments were carried out in accordance with relevant guidelines and regulations. The protocols were approved by the University of Camerino, by the Applied and Environmental Botany department.

The animals were tagged with nylon harnesses, according to standard procedures [49, 50]. They appeared to tolerate the tag attachments well, based on reports from the farmers and the fact that no anomalies to fur or feathers were found when retrieving the tags. We recorded the 3D-acceleration of the tagged animals continuously at 54Hz during the Oct-Nov 2016 period, and at 54Hz every 120 seconds for 3.5 seconds during the Jan–Apr 2017 period. We calculated the ODBA according to standard procedures [51–53]. The two dogs (of 4 on the farm) were initially restricted to a narrow farm yard, but later roamed the pastures with the sheep. A total of ca. 20 cows were chained next to each other inside a stable during Oct 2016 to Mar 10^th^ 2017, but were free to roam on a pasture after Mar 11^th^ 2017. The sheep were kept free-running inside a stable (ca. 4 by 20 meters) in a group of about 100 animals from Oct 2016 to Mar 10^th^ 2017. Later the same group was kept free-running in open pastures.

### Data Description

We used 3D acceleration sensors to measure the activity of the animals (Fig. 3C). As only cows, dogs and sheep were available in all three time periods of the study (Oct-Nov 2016, Jan-Mar 2017 and Mar-Apr 2017), only these three species were considered in the analysis. For each of these three species, we computed the 15 min average of their ODBA, i.e. the average acceleration and the average over all tagged animals of the respective species.

**Fig. 3.**
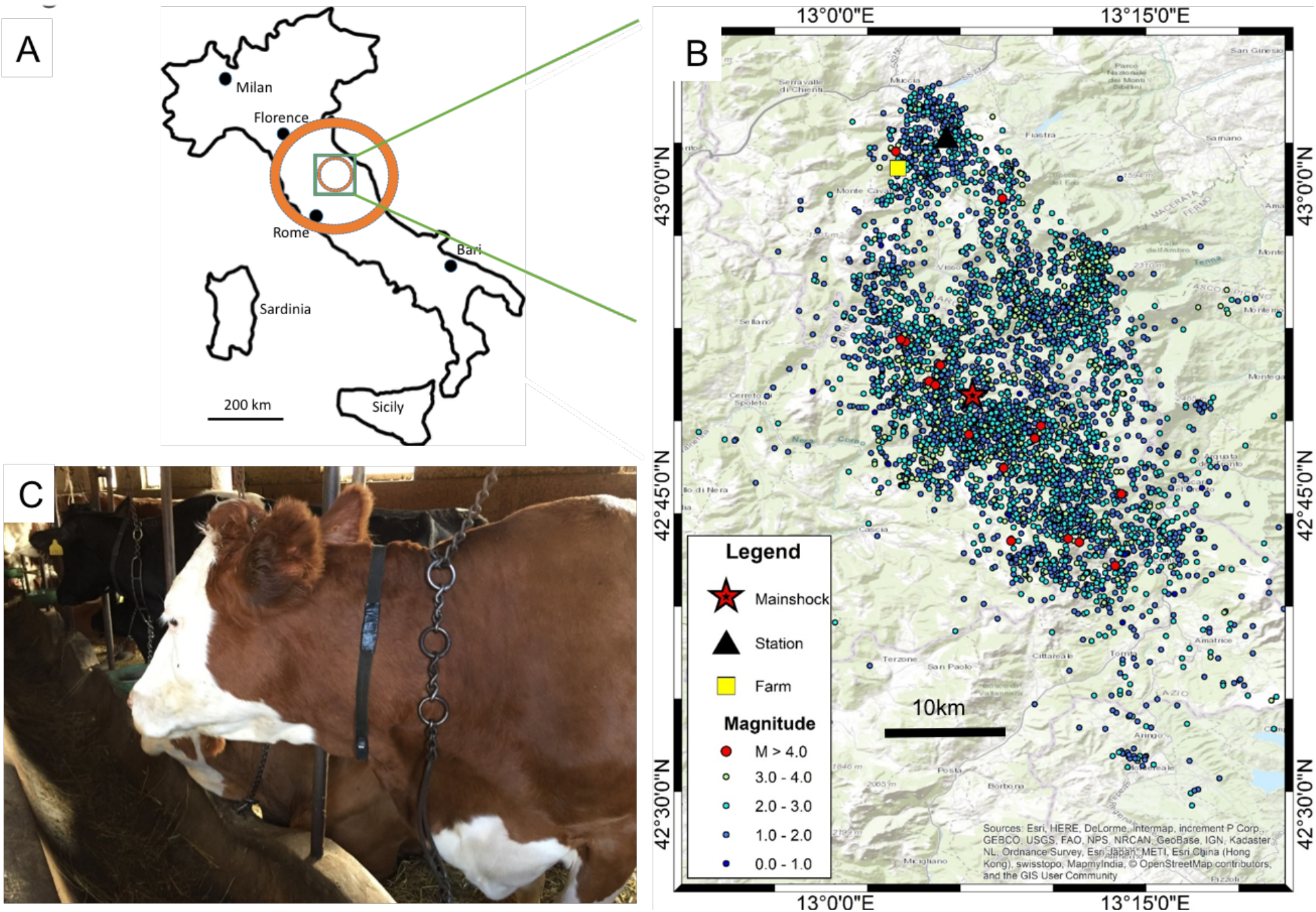
A) Map showing the locations of the Oct-Nov 2016 earthquakes in central Italy. Concentric circles show the area where the impact of the earthquake was perceived by people. The largest circle represents the M6.6 earthquake from Oct 30, 2016 at 6:40 UTC. Data from www.earthquaketrack.com, based on original data from https://earthquake.usgs.gov/. B) Detailed map showing the epicenters of the earthquakes studied during the current investigation, as well as the location of the farm where animals were studied. C) Example of a tagged cow inside a stable secured next to other cows. Electronic tag sits ventral on black neckband.

Between Oct 29^th^ and Nov 7^th^ 2016, the animals experienced a total of 5304 earthquakes with M > 0.4 (maximum M 6.6) and from Jan to Apr 2017 a total of 12948 (maximum M 4.2, Fig. 4,5,6). The M6.6. Norcia mainshock was felt throughout central Italy and into Rome (Fig. 3A).

**Fig. 4.**
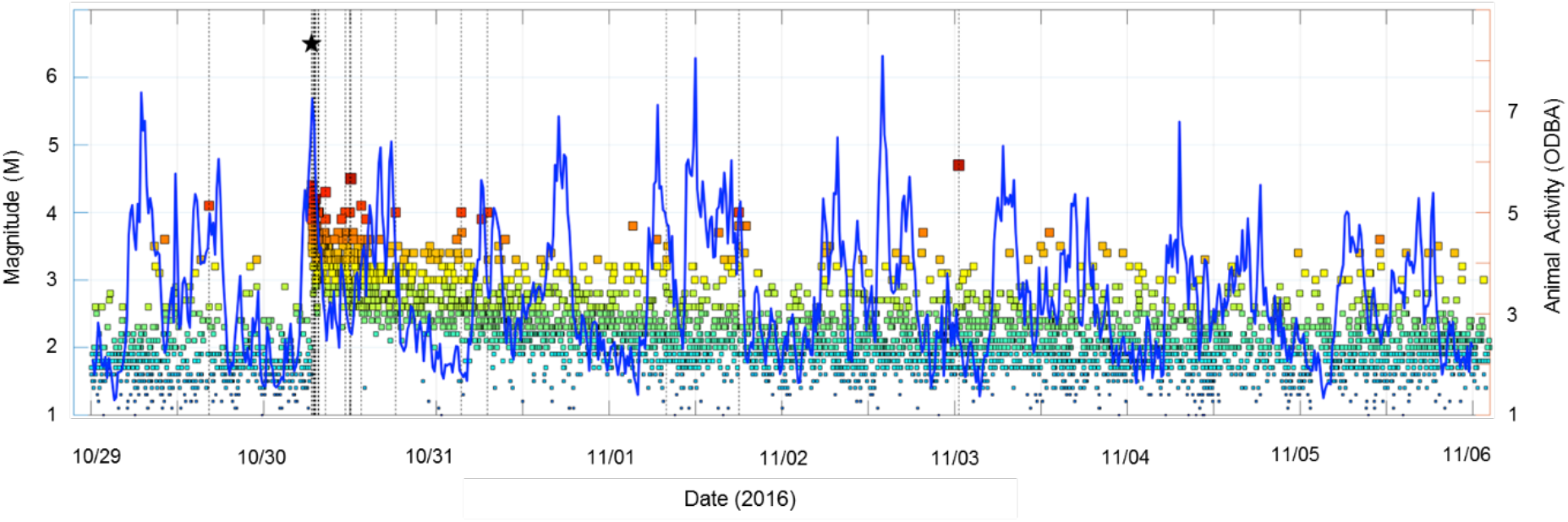
Animal activity in relation to earthquake activity. The blue line depicts the ODBA-value summed over 15min for farm animals during the period Oct 29^th^ – Nov 6^th^ 2016. Colored symbols mark earthquake activity in the region (increasing with redness indicates higher magnitudes) dots showing earthquakes above M4, and a black star indicating the M6.6 Norcia earthquake of Oct 30^th^ 2016.

For each of the earthquakes, we had information on the magnitude and time, but also the latitude, longitude and depth of the hypocenter, which allowed us to compute the hypocentral distance between the farm and the respective earthquakes. The hypocentral distances of these earthquakes range from 5 to 28 km to the tagged animals, oriented mostly in a southerly direction (Fig. 3B).

Fig. 4 shows the time series of both, aggregated animal activity (average over the species) and the earthquakes for the Oct-Nov 2016 period.

### Data Preprocessing

Regarding the animal activity, we could directly use the ODBA time series (on 15 min intervals) of the three species during the three periods.

Regarding the earthquake activity, a measure for the actual earthquake activity at the farm was needed. The magnitude is a measure for the strength of the earthquake that does not consider the distance from the hypocenter. As measure for earthquake activity at the farm, we estimated the peak ground acceleration (PGA) at the farm for the entire available earthquake catalog (0.4 < M < 6.6) using a regional ground-motion prediction equation [54](S3).

The depth-dependent seismic wave speeds in the upper Earth crust range typically from 3-7 km/s for the primary P-waves, and 2 - 4.5 km/s for the secondary S-waves. Given that we measure animal activity in 15 min time intervals, the travel times of seismic waves from the earthquake’s hypocenters and the farm is on the order of a few seconds, and hence negligible in our analysis

Fig. 5 visualizes the spatial distribution of the earthquakes, color-coding each earthquake event by the respective PGA at the farm.

**Fig. 5.**
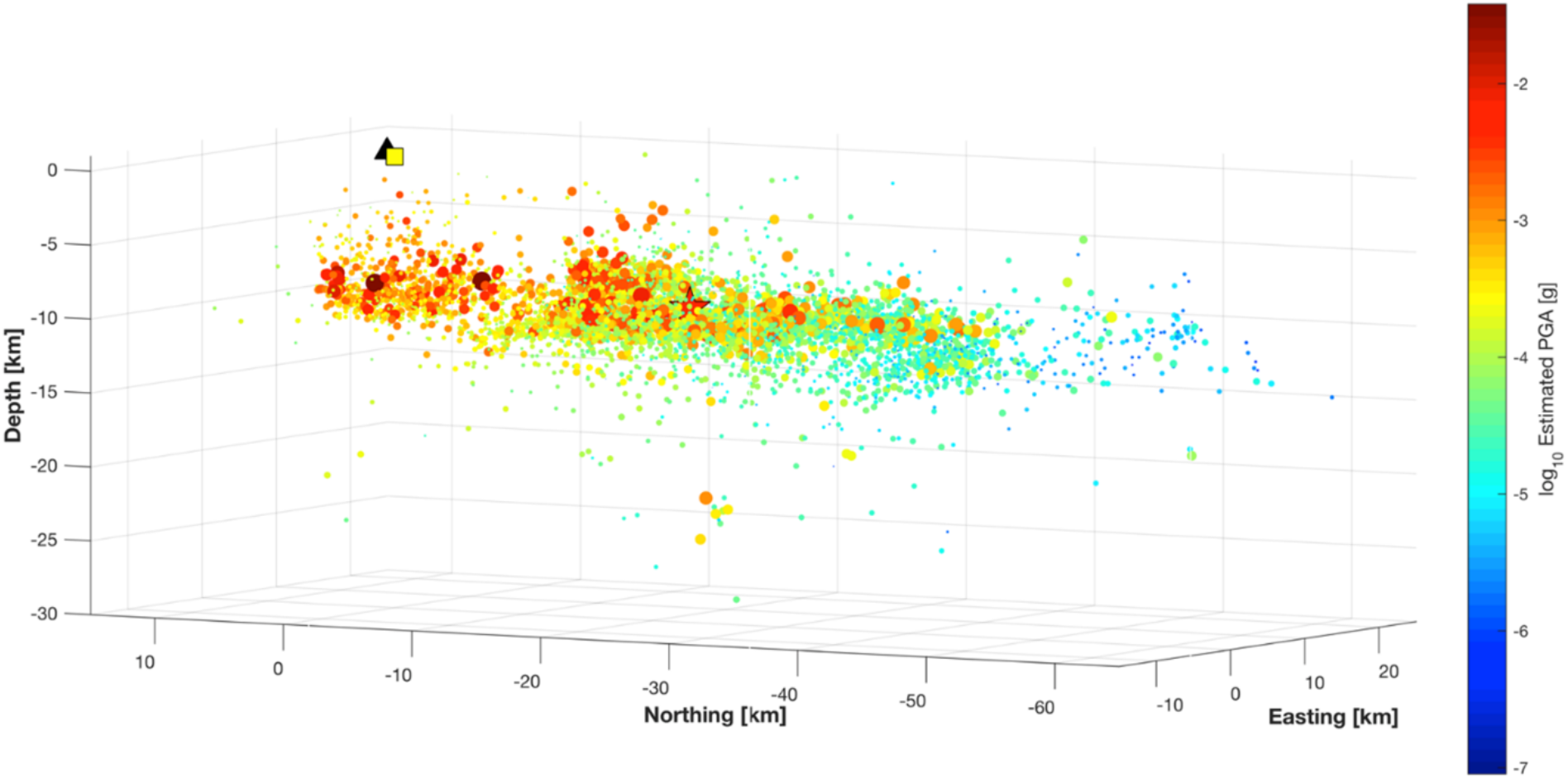
Spatial distribution of earthquakes during the Oct 29^th^ – Nov 6^th^ 2016 period. Colored dots show estimated peak-ground-acceleration (PGA) at the farm (black triangle) for each event, computed using the empirical relations of Bindi et al. (1), considering magnitude, hypocentral distance, faulting style and soil-site class (here class A, with VS30 = 800 m/s (Lucia Luzi, pers. comm.)). Larger far-distant events may cause stronger shaking then nearby small events [units in log10(g), g = 9.81 m/s2; the Norcia main shock was about 0.1 g at the farm)].

The time series of estimated PGA events at the farm are depicted for Oct-Nov 2016 and Jan-Apr 2017 in Fig. 6A and 6B, respectively.

**Fig. 6.**
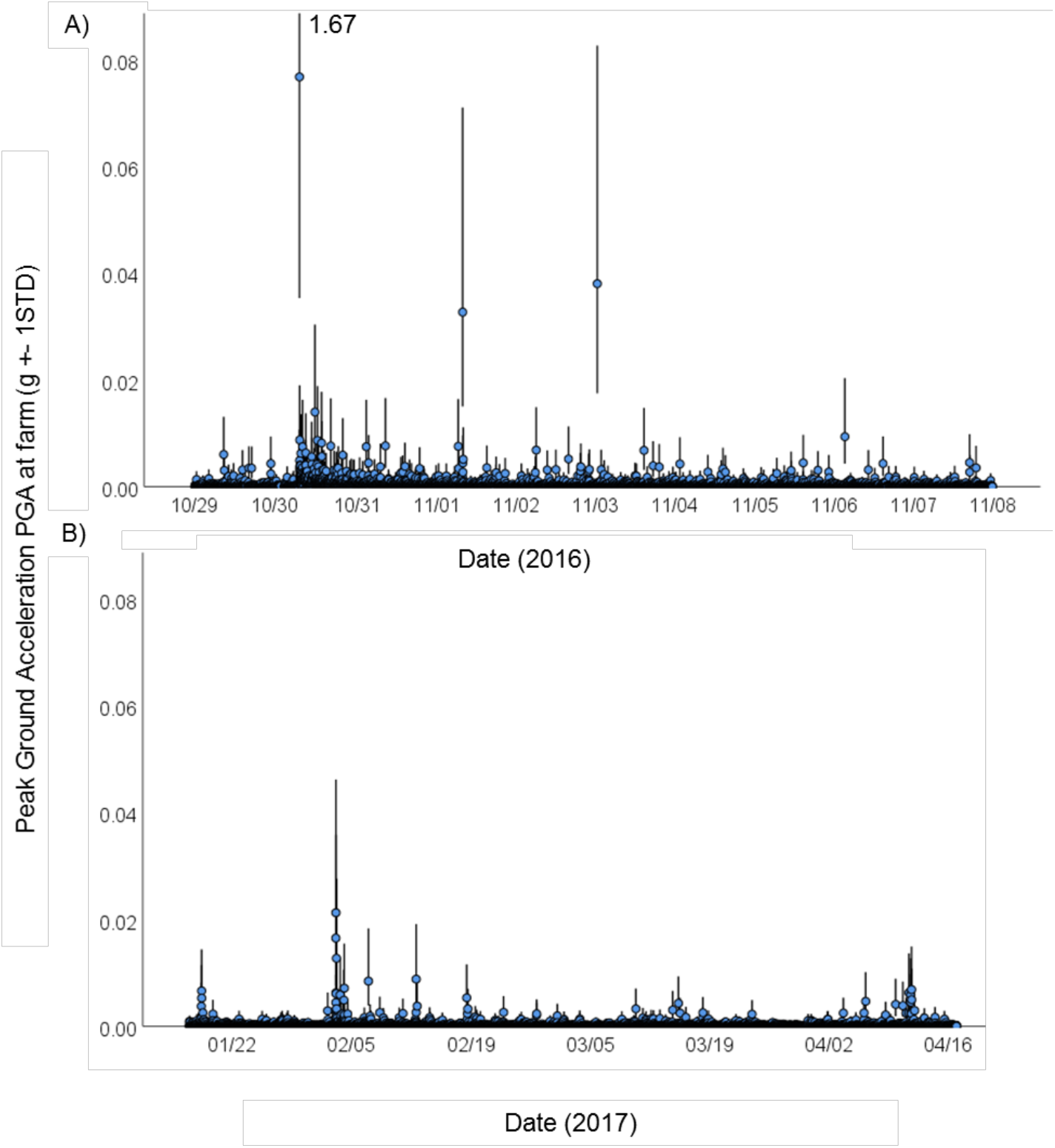
Time series of peak-ground-acceleration (PGA) at the farm, during A) the Oct 29 – Nov 06, 2016 and B), the Jan 17 – Apr 17, 2017 period. Blue dots indicate estimated PGA for each event as in Fig. 4,5. Vertical lines highlight +− 1 STD of the estimates. Please note the different x-axis time scales, and the upper truncation of the highest PGA value (for graphical comparability between A and B).

To align the time-dependent earthquake-specific information with the animal activity data, we consider an aggregated measure of the estimated PGA-value for each earthquake, using the highest PGA-value estimated for any event that occurred in the respective time span. Note that the results of the study are robust against the choice of the aggregation method (alternatives were the mean and the sum of PGA events in the respective window).

After this processing, we obtain time series of animal activities and earthquake measures sampled at identical time intervals.

### Retrieving the daily patterns in animal behavior

The animal activity time series show daily patterns. We estimated the daily patterns separately for the three animal species and the three periods, using Fourier series with 24 hours periodicity. 15 frequencies (15 sin, 15 cos and one constant term) were the largest number of frequencies selected by the Bayesian information criterion (BIC) for any of the time series, so we used 15 frequencies for all series for comparative purposes. However, the findings of our analysis are robust against the chosen number of frequencies used for the estimation of daily patterns (we tested 10-25 frequencies).

### Vector Autoregressive (VAR) Analysis

In a vector autoregressive model, the variable vector (here the ODBA values of the three animal groups and the PGA activity) at time t depends linearly on past (lagged) values of the variable vector at times t-1, t-2, …, t-p, where p denotes the number of lags. Using this analysis allows to assess the mutual influence of the variables on each other, e.g. via Impulse Response Functions.

Based on the excess animal activity (observed activity – estimated daily activity) of cows, dogs and sheep and the PGA time series, we estimated vector autoregressive processes for the three periods. The maximum number of lags suggested by BIC for any of the three periods was 6 (VAR(6)), so we used 6 lags in all three periods for comparative purposes. However, the results are robust with respect to the number of lags used in the VAR model (we considered 4-10 lags).

The VAR model and the resulting Impulse Response Functions (IRFs) show the mutual influences of the animal species on each other as well as the influence of earthquakes on the animal species (Fig. S7, S8). The mutual influence of the animal species and the reaction to earthquakes is part of the normal behavior of the animals. To obtain the abnormal animal activity, in which we want to find anticipatory patterns, we subtract both, daily patterns and the predictions of the VAR model from the observed animal activities.

### Threshold Analysis

Consider the abnormal component of the animal activity, as defined in the previous paragraph, i.e., abnormal activity = observed activity – daily patterns – VAR predictions. The following steps were applied for each of the animal species, as well as for the aggregate animal behavior (mean of animal species) in all three periods.

At first, each PGA event that exceeded a given threshold (e.g. 1.5 standard deviations above the mean) was selected. For each of these PGA events, we selected all occurrences of ‘unusual’ animal activity in a time span up to 20 hours before the respective PGA event that exceeded a second threshold (e.g. 2.5 standard deviations above average ‘unusual’ animal activity). In this way, we created pairs of unusually high PGA and animal activities.

For each pair of observations found in this manner, we compute the respective anticipation time (time of PGA event – time of abnormal animal activity event) and plot it against the hypocentral distance between the farm and the respective earthquake event.

The results are robust with respect to the choice of the thresholds (Fig. S10) and with respect to outliers due to the use of median regressions.

#### Informed consent

We received informed consent of all study participants for publication of identifying information/images in an online open-access publication.

## Acknowledgments

We are indebted to the Angeli family for the help, support and understanding despite their personal hardships. P.M.M. acknowledges L. Parisi and L. Lombardo (both at KAUST) for their help in processing the earthquake data collected by the INGV Rome (Istituto Nazionale di Geofisica and Volcanologica). This study was funded by the Max Planck Society and partially by King Abdullah University of Science and Technology (KAUST), BAS/1/1339-01-01.

## Author contributions

M.W., U.M., M.B., L.D. and D.K. developed the concept, M.W., U.M., initiated the study and collected the animal data, P.M.M. contributed data and analysis, W.P., G.F. and M.W. analyzed the data and all authors contributed to the discussion and final write-up process.

## Additional information

**Supplementary Information** accompanies this manuscript.

## Supplementary materials

### 1. Preparation of equipment and selection of field site and animals

Starting in January 2014, we kept 20 GPS and 3D-acceleration data loggers (22 grams, E-Obs, Munich, Germany) in a state of constant readiness for a potential rapid use on animals in earthquake areas. The rechargeable batteries of these data loggers were kept at high voltage through a recharge every month. The 3D-acceleration sensors were initially calibrated according to standard procedures (*1–3*). We also kept various sizes of nylon harness material in stock that could be adjusted to previously unknown sizes and shapes of the animals we hoped to tag and observe.

### 2. Analysis: The daily patterns of animal activity

We found that the three animal species showed different daily activity patterns in the stable and on the pasture. In the stable (Oct-Nov 2016 data and Jan-Mar 10, 2017 data), animals were active in the morning and afternoon, but at mid-day they were relatively calm (see Fig. S4). However, dogs and sheep showed no longer a reduced activity around noon when roaming freely on the pastures. Moreover, the volatility of cows and sheep was small while in the stable, but on the pasture both dogs and sheep showed rather volatile behavior, thus deviated often and strongly from the daily patterns.

### 3. Analysis: Earthquake peak ground acceleration (PGA)

The effects of the earthquake activity on the farm animals was estimated by calculating the expected horizontal peak ground acceleration (PGA) at the farm for each earthquake in the catalog. This was accomplished by applying the empirical ground-motion prediction equation (GMPE) (*10*) using the corresponding earthquake magnitude and the hypocentral distance to the farm. The correct time of the respective activities at the farm was calculated using the distance between the hypocenter and the farm and an depth-averaged seismic P-wave speed of 6.15 km/s, based on published Earth-structure models for the region (*11, 12*).

To match the timescale of the animal ODBA data, we analyzed PGA-values of events that occurred in the same 15min timescale, using two different approaches. The first method used the maximum peak ground acceleration (PGA) at the farm of all earthquakes that happened during the respective 15 min intervals; the second method is based on the sum of observed PGA values in the respective 15 min interval. The subsequent analysis is based on the maximum PGA value during each 15 min interval. However, the results are robust against the choice of the PGA-accumulation method.

### 4. Analysis: Vector Autoregressive (VAR) estimation (mutual behavior of animals and reactive behavior after earthquakes)

The Impulse Response Functions (IRF’s) show that the different animal species reacted significantly upon each other. However, the interaction patterns differed between the periods in the stable (Oct-Nov 2016 and Jan-Mar 2017) and the period on the pasture (Mar-Apr 2017). In the stable (see Fig. S7), the cows’ activity significantly increased after a shock in the dogs’ activity, leading to a significantly increased activity for about 1 hour, followed by a calming down phase (row 1, column 2). Dogs instantaneously increased activity after a shock in the cows’ activity, but very quickly relaxed afterwards (row 2, column 1). On the pasture (see Fig. S8), the patterns as described above remained valid, only sheep now strongly reacted on the dogs’ activity (row 3, column 2), most probably since dogs were used to guard sheep on the pasture.

The time needed to calm down after a shock differed between animal species. In the stable or on the courtyard, dogs were back to normal activity after about 30 min, cows after about 45 min and sheep after about one hour (see Fig. S7, plots on the diagonal). This order remained unchanged on the pasture, but the animals needed in general more time to calm down.

In terms of the animals’ reactive behavior on earthquakes, we found that dogs were most sensitive, followed by cows. Sheep seemed to be least sensitive. For the Jan-Mar 2017 data (animals in the stable), we found significant reactions of dogs and cows on the earthquakes (see Fig. S7, row 1, column 4 and row 2, column 4), whereas the activity of the sheep showed no significant response. The functional form of the dogs’ response was confirmed by the Oct-Nov, 2016 data. For the Mar-Apr 2017 data (animals on the pasture), no animal group reacted statistically significantly on the earthquakes (see Fig. S8). Possible explanations are the higher noise on the pasture and that animals might be less afraid during earthquakes when they are roaming freely outside the stable.

Moreover, we found that dogs and cows reacted differently on earthquakes. Whereas the dogs’ activity increased after earthquakes and faded into a relaxing period later on (see Fig. S8, row 2, column 4), the activity of cows first remained constant or even slightly decreased, before it increased significantly and finally faded. Knowing that cows reacted strongly upon dog behavior, we expected that the cows’ post-earthquake activity in the absence of dogs would decrease, thus an earthquake would calm them down instead of increase their activity. However, due to the increased activity of the dogs, we found that, after the first period where the influences of the earthquake and the dogs were balanced, the activity of cows increased due to the dogs’ increased activity. To confirm this experimentally, one could observe cows on farms where they are separated from the dogs.

We conclude that different animal species behaved differently during earthquakes and that their mutual interactions were significant and relevant to describe their behavior. To detect the animals’ specific reactions towards earthquakes more accurately, future research designs should aim to remove (or at least mitigate) the impact of possible interdependencies between the animal species. Furthermore, significant responses of different animal species on earthquakes were only visible when the animals were restrained, i.e., in the stable (sheep and cows) or in a narrow farmyard (dogs).

These results were all robust against changing the number of lags in the VAR estimations.

### 5. Analysis: Threshold Model (Anticipatory Animal Behavior Prior to Earthquakes)

In this Threshold Model, we selected PGA occurrences that exceeded a given threshold and then searched for animal activity beyond a second threshold in a 20h time window before this threshold-exceeding earthquake event. For all pairs of PGA and animal activity found in this manner, we plotted the time difference between the PGA occurrence and animal activity (warning time) against the hypocentral distance of the farm.

Via the thresholds, we filtered the relevant data for estimating the relationship between warning time and distance. Thresholds that were too high led to very few observations; too low thresholds resulted in increased irrelevant noise. For the animal activity we considered thresholds in the range of 2 to 3 standard deviations, for the earthquakes we considered thresholds in the range of 1 to 2 standard deviations. We considered smaller thresholds for the earthquake than for the animal data, since the empirical distribution of the PGA events was more centered around zero than the distribution of the abnormal animal activity.

For the selected data pairs, we estimated a linear relationship between warning time and hypocentral distance by ordinary least squares (OLS) and median regression. Considering the aggregated animal activity, we obtained significantly negative slopes for the Oct-Nov 2016 and the Jan-Mar 2017 periods (animals in the stable) (see Fig. S9). Significance was absent for the Mar-Apr 2017 period on the pasture (See Fig. 1), equivalent to the absence of significant reactive behavior on the pasture in the VAR analysis.

We obtained the best (in terms of statistical significance) regression results for the aggregated animal data, which smoothened the activity levels of the various animal species and therefore helped to reduce noise in the abnormal activity. Considering the data on the animal species separately resulted in statistically insignificant slope parameters.

The results are robust against the threshold choices (see Fig. S10) and the estimation method for the linear function. Moreover, we varied the time window before a threshold-exceeding earthquake in which we searched for threshold-exceeding animal activity between 15 hours and 25 hours and found the results to be robust. Whenever we chose search windows, however, that were too narrow in time, they possibly cut out important observations for earthquakes in close spatial proximity and their associated potential long animal warning times. On the other hand, time windows that were too long might contain too many irrelevant observations.

## Supplementary Figures and Tables

**Figure S1.**
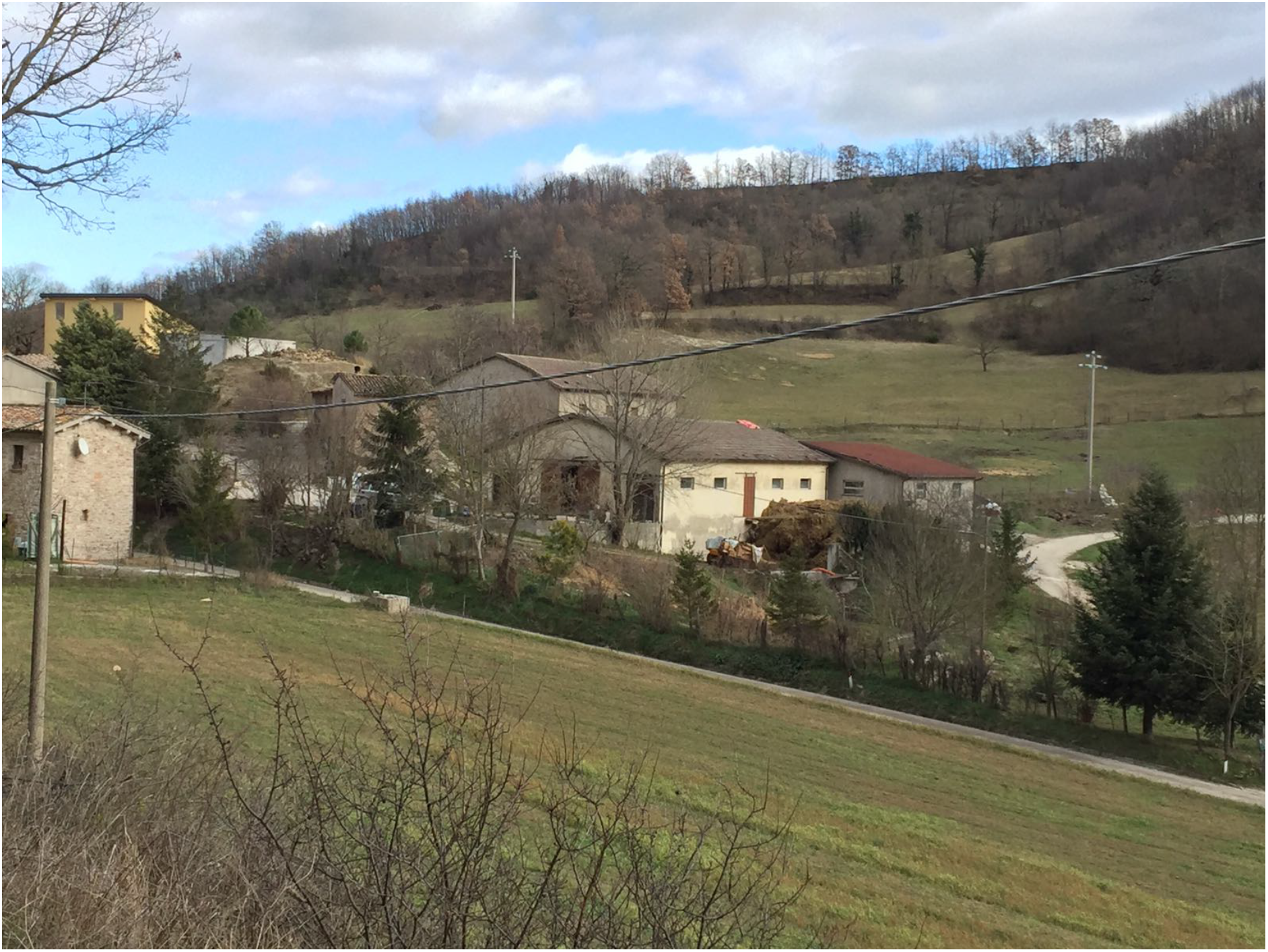
The Angeli brothers farm in Capriglia/Italy and its surroundings. The stable of the cows is depicted in the middle of the picture.

**Figure S2.**
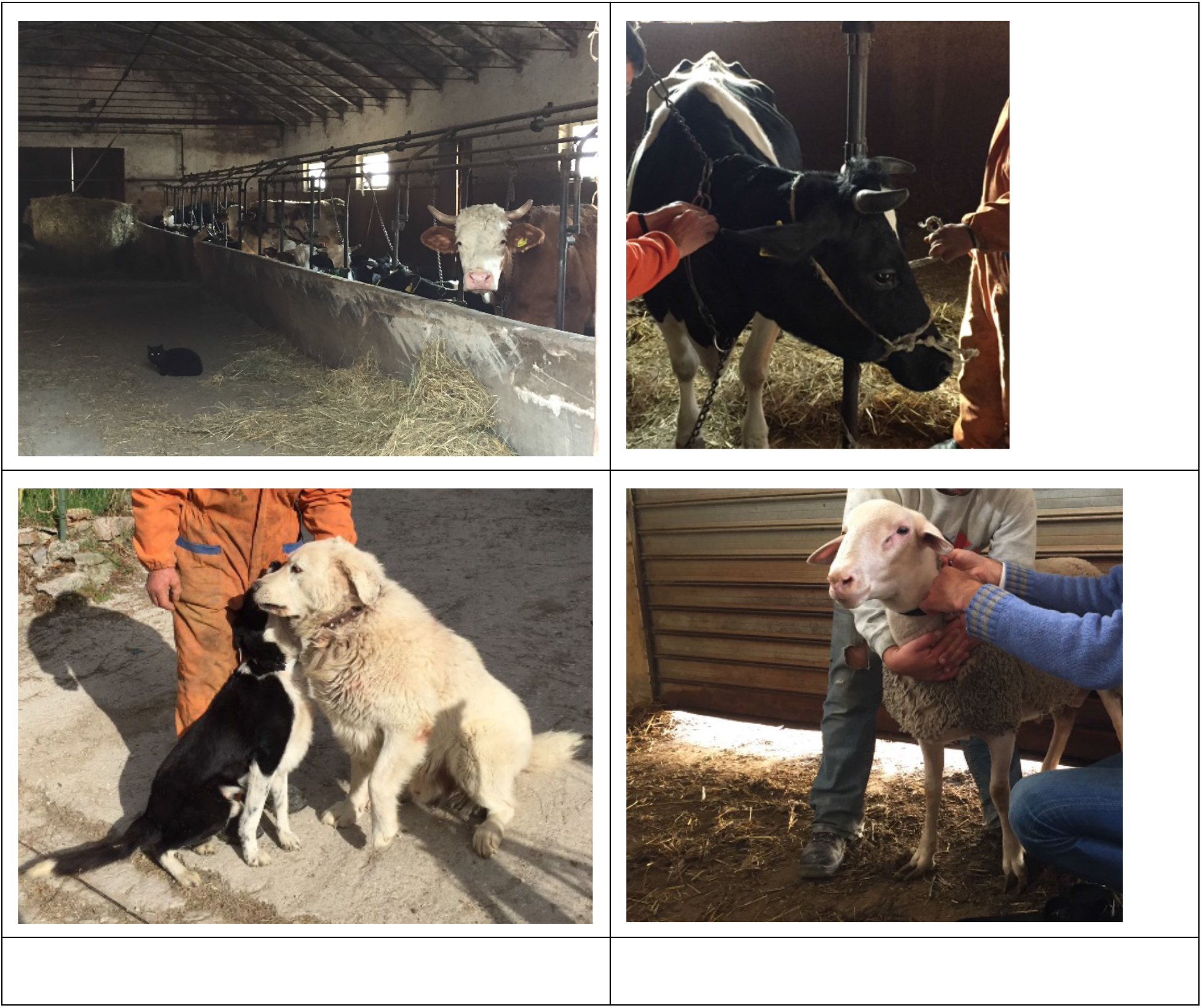
Photos showing the tagged animals during the winter (Oct 2016 – Mar 10, 2017) period. Top left: Cows lined up in the stable. Top right: Tagging of individual cow. Bottom left: Dogs in the farm yard. Bottom right: Sheep in the stable.

**Figure S3.**
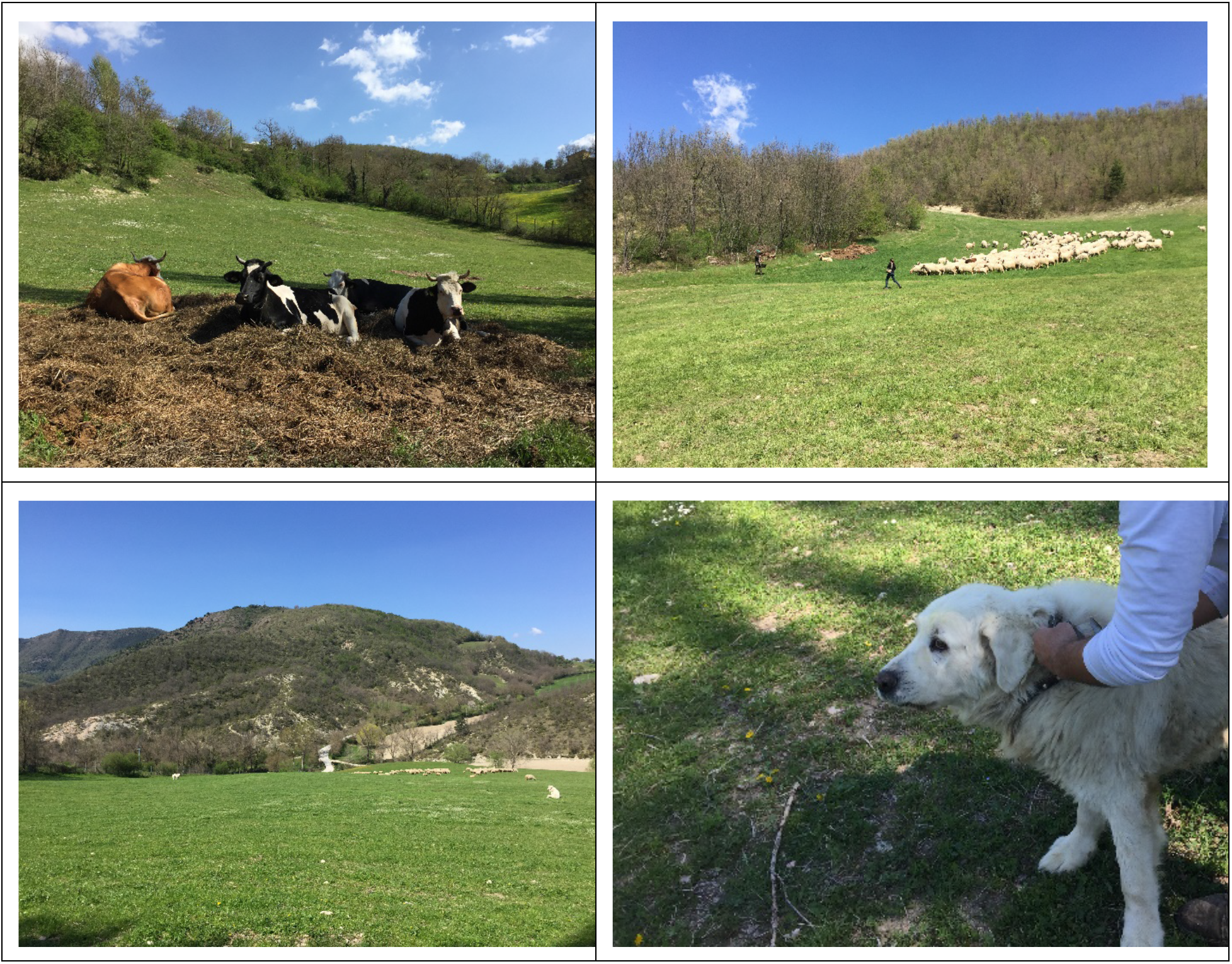
Photos showing the tagged animals during the spring (after Mar 11, 2017) period. Top left: Cows on the pasture. Top right: Sheep on the pasture. Bottom left: Dog on the pastures guarding sheep. Bottom right: Dogs on the pasture.

**Figure S4.**
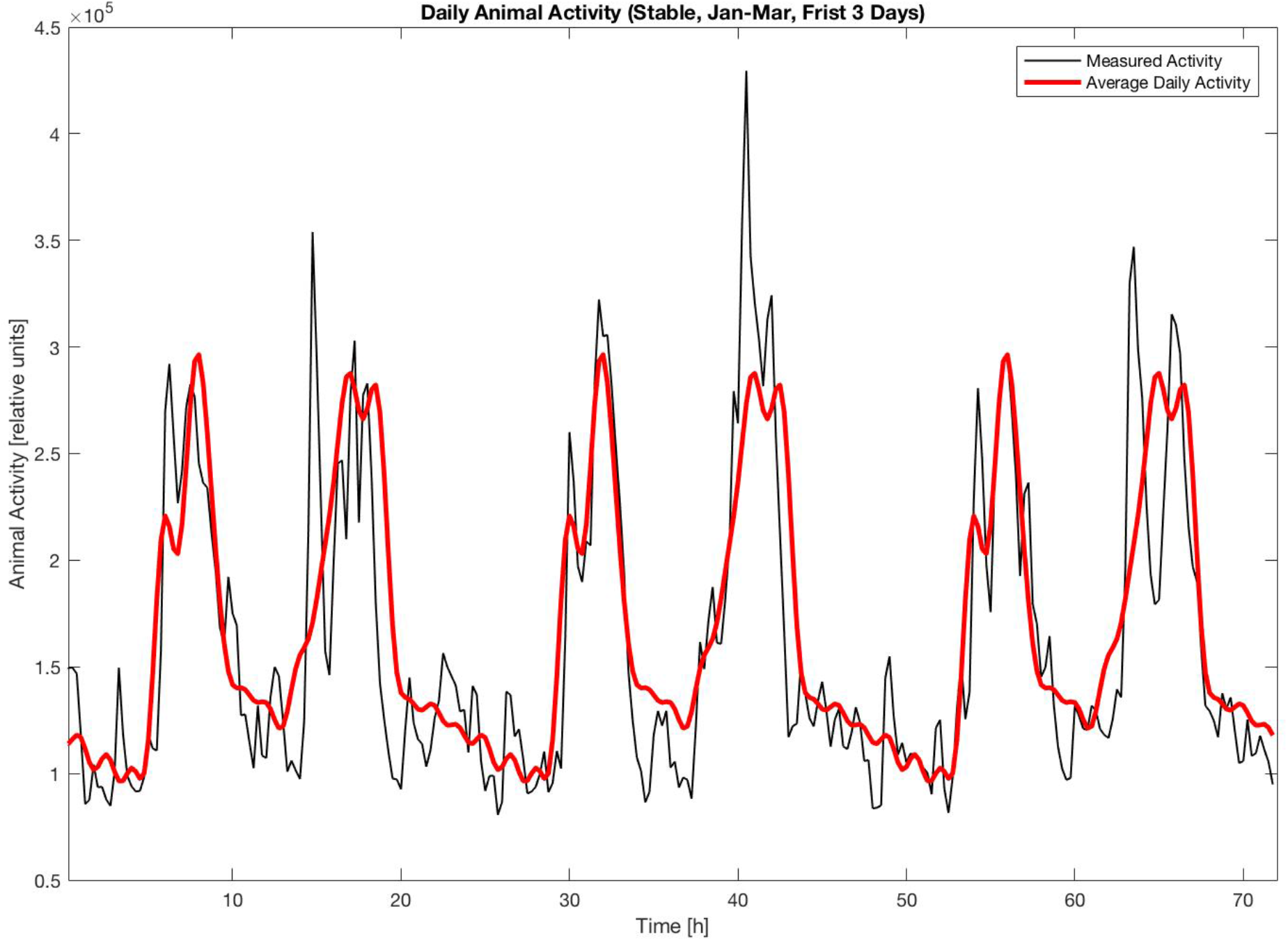
Aggregated animal behavior and estimated daily patterns. The animal activity is plotted for the first three days of the Jan-Mar 2017 data (animals in the stable). The actual animals’ behavior is shown in black, the estimated daily patterns are given in red. The animals show high activity during the morning and afternoon, but are relatively calm during noon. The animal activity is measured by the accumulated ODBA in relative units.

**Figure S5.**
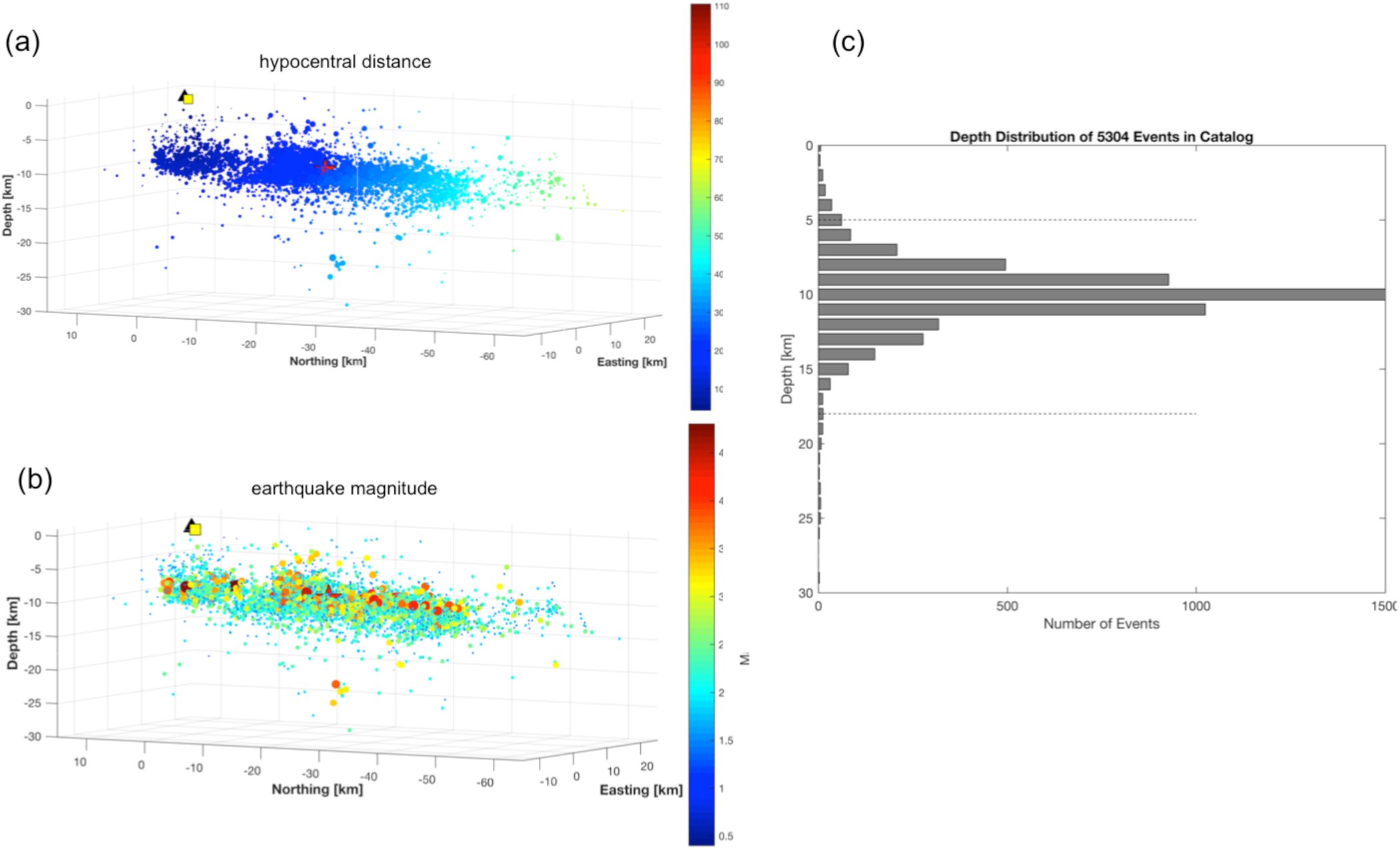
Earthquake catalog. Properties of the earthquake catalog for the time period 28/10/2016 – 08/11/2016, consisting of 5304 earthquakes in the magnitude range 0.4 ≤ M ≤ 6.5. (a) 3D view of earthquake locations with color-coded hypocentral distances. The yellow square marks the position of the farm, to which the spatial data are referenced. (b) Same as in (a), but color-coded by magnitude. Notice that symbol-size is also magnitude-dependent. (c) Depth distributions of all hypocenters. 95% of hypocenters fall in the depth range of 5-18 km (indicated by the two horizontal lines). Note that c) is similar to Fig. 5 of the main part.

**Figure S6.**
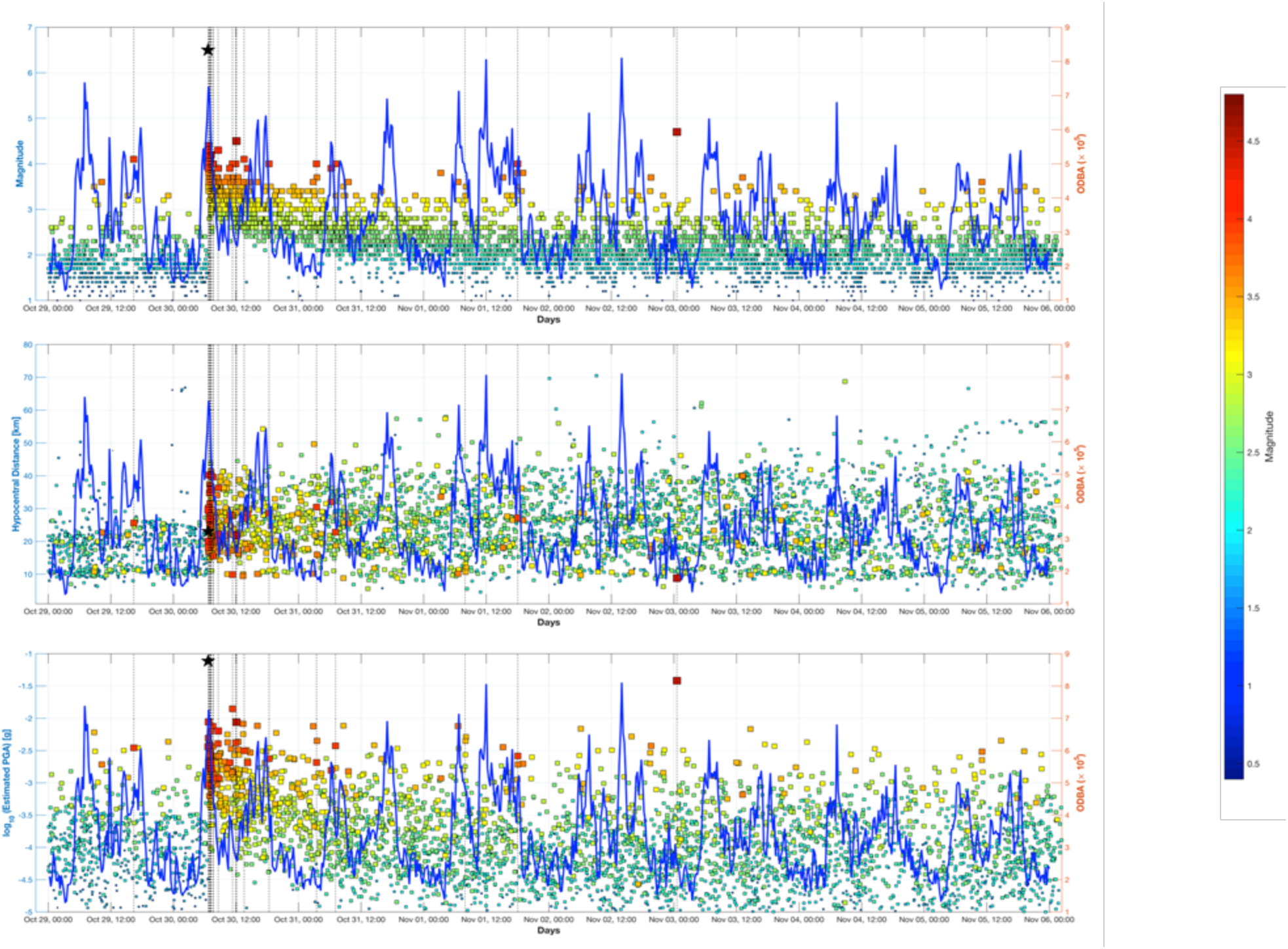
Comparison of ODBA measurements with earthquake data. Temporal analysis of earthquake data: top – earthquake magnitude; center – hypocentral distance from the farm; bottom – estimated peak ground acceleration (PGA). ODBA shown in blue; earthquake magnitude is color-coded; M ≥ 4 also marked by vertical lines. PGA is measured in multiples of the gravitational constant g (= 9.81 m/s^2^).

**Figure S7.**
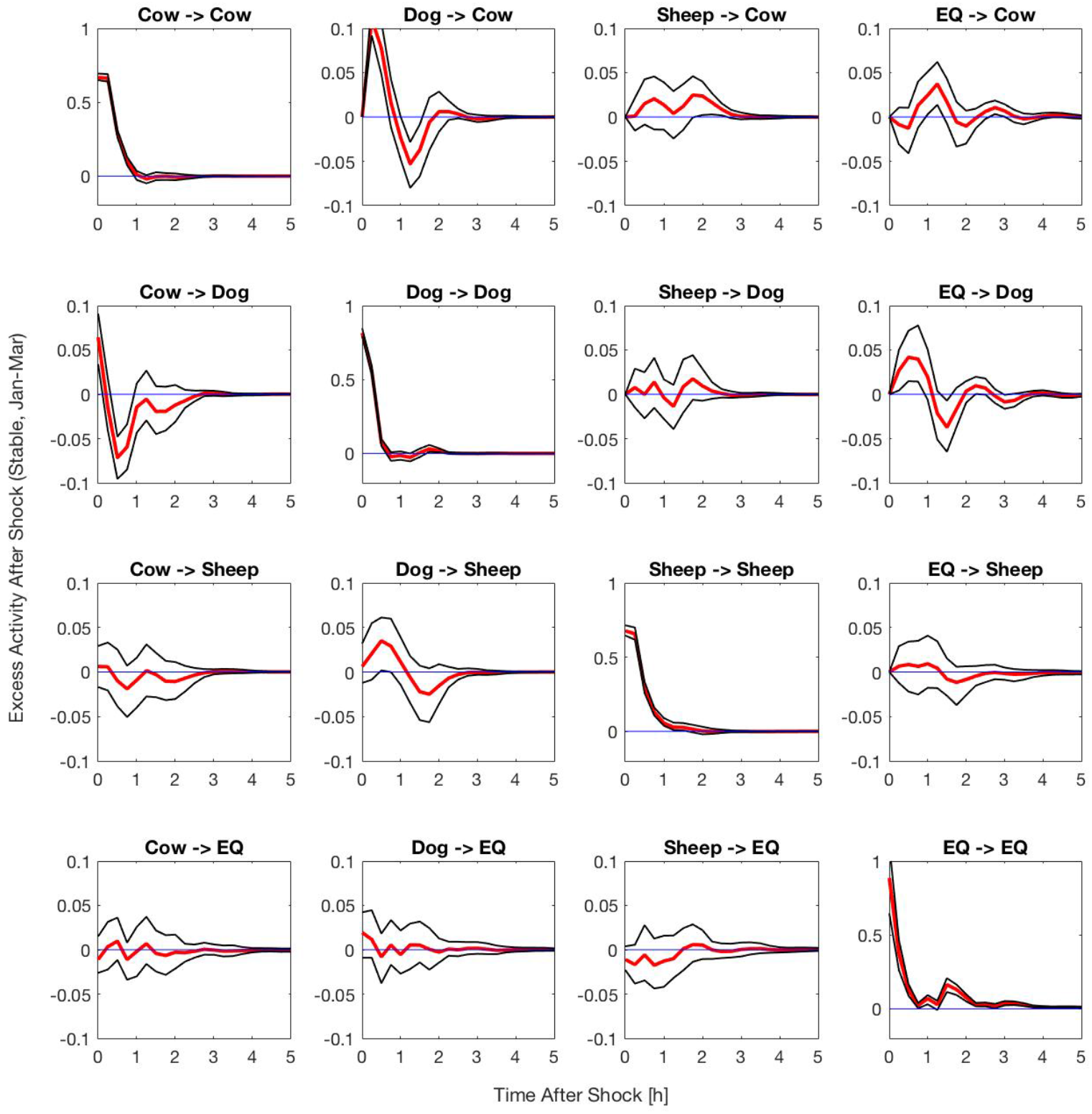
How animals react to earthquake in a building: Impulse Response Functions (IRF) in the stable. The plots show the statistical reaction patterns of the animal species and the PGA on orthogonalized shocks in each of the time series for the Jan-Mar 2017 data in the stable, in relation to the time after the shock. The red line shows the VAR orthogonal impulse responses, the black lines the ±95% confidence interval (CI). Each plot examines the effect of one variable on another one (described above the plot, e.g., the effect of abnormal dogs’ activity on abnormal cows’ activity, termed Dog -> Cow’). Whenever the 95% CI does not overlap with zero line, there is a statistically significant influence. The different animal species reacted upon activity shocks of the other animal species and on earthquake occurrence. While dogs showed increased activity after an earthquake (row 2, column 4), cows seemed to be rather unusually calm (row 1, column 4) and were then influenced by the increased activity of the dogs (row 2, column 1).

**Figure S8.**
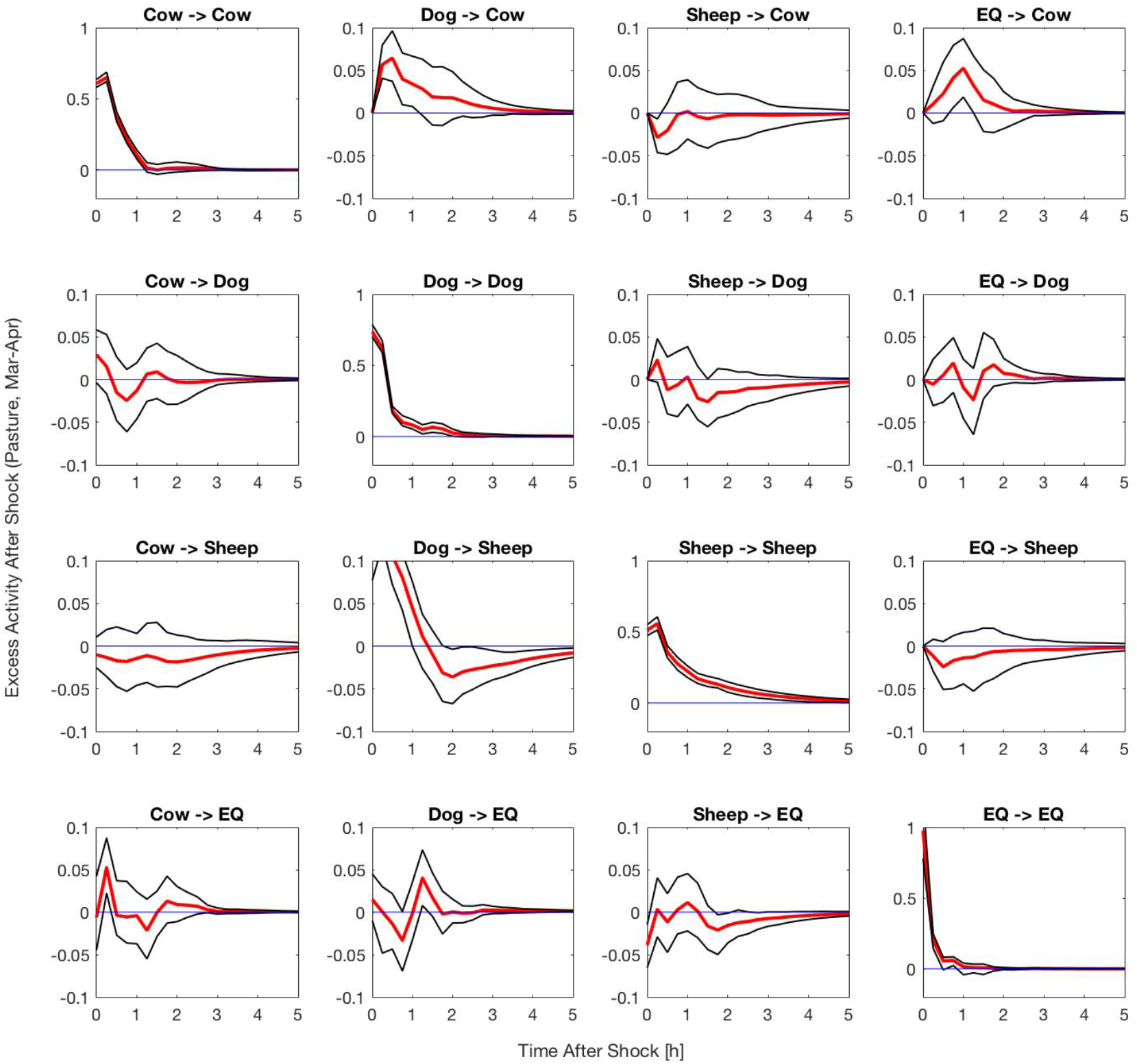
How animals react to earthquake on a pasture: Impulse Response Functions (IRF) on the pasture. The plots show the statistical reaction patterns of the different animal species and the PGA on orthogonalized shocks in each of the time series for the Mar-Apr 2017 data on the pasture. The animals still reacted upon each other, but no significant reaction on PGA shocks is visible. However, sheep reacted significantly on dog activity, most probably since the dogs guarded the sheep on the pasture, but not in the stable. Please see detailed description of Fig. S7.

**Figure S9.**
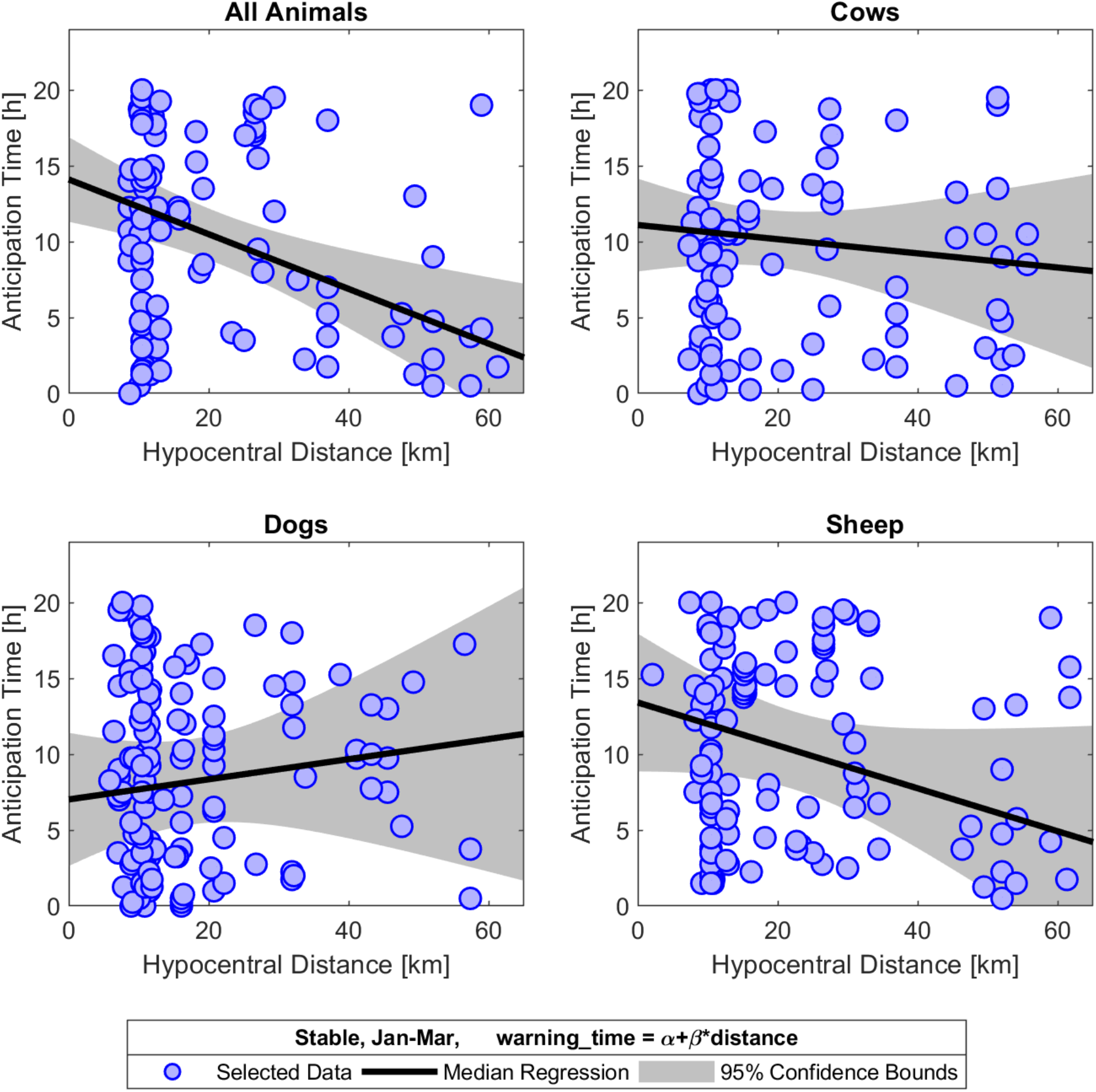
How animals anticipate an earthquake: Distance – Warning Time Plots and Estimated Linear Relationship. For the Jan-Mar 2017 data in the stable, we plotted the computed animal warning time (see main text) against the distance between earthquake hypocenter and farm. We estimated a linear relationship (black line) by median regressions (OLS results are equivalent) and also provide the 95% confidence bounds (gray area). Whereas the relationship for the separate animal species is statistically insignificant, we find a significantly negative relationship for the aggregated animal data.

**Figure S10.**
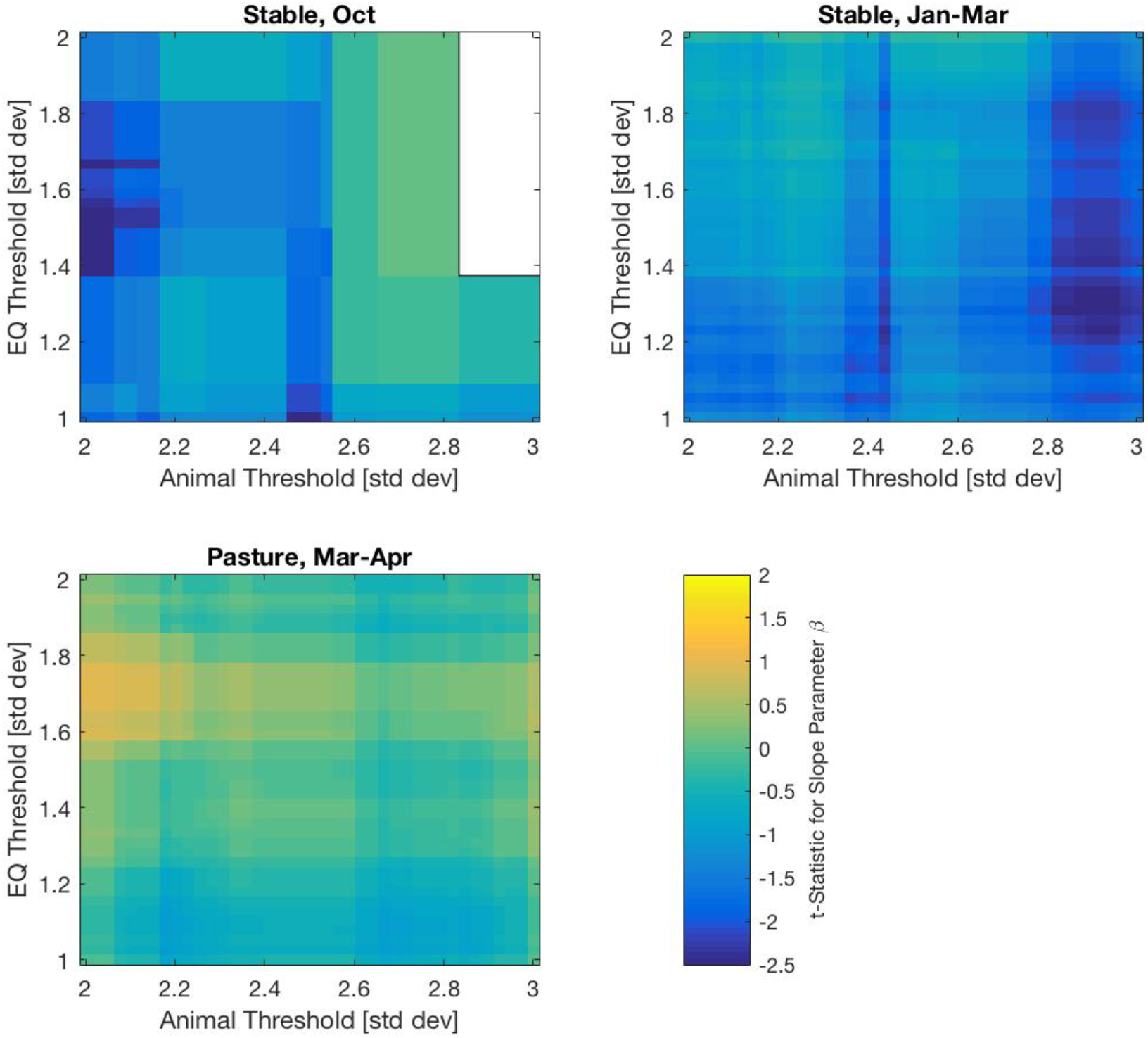
Slope T-Statistics and Threshold Choices. The heatmaps show the *t*-statistic for the estimated slope parameters of a linear relationship. If the t-statistic is above 1.96, the slope parameter is significantly larger than zero, if the t-statistic is below 1.96, the slope parameter is significantly lower than zero. We found threshold combinations that yielded a significantly negative relationship between warning time and distance for both time periods, i.e., Oct – Nov, 2016 and Jan 17 – Mar 10, 2017, when the animals were in the stable (‘Jan-Apr Stable’), but not for the period when they were on the pasture (Mar 11 – Apr 14, 2017, termed ‘Jan-Apr Pasture’).

